# Monoclonal antibody stability can be usefully monitored using the excitation-energy-dependent fluorescence edge-shift

**DOI:** 10.1101/2020.03.23.003608

**Authors:** MK Knight, RE Woolley, A Kwok, S Parsons, HBL Jones, CE Gulácsy, P Phaal, O Kassaar, K Dawkins, E Rodriguez, A Marques, L Bowsher, SA Wells, A Watts, JMH van den Elsen, A Turner, J O’Hara, CR Pudney

**Affiliations:** UCB, 216 Bath Road, Slough SL1 3WE, United Kingdom; Department of Biology and Biochemistry, University of Bath, Bath, UK; Department of Physics, University of Bath, Bath, UK; Department of Pharmacy and Pharmacology, University of Bath, Bath, UK; Centre for therapeutic Innovation, University of Bath, Bath, UK

**Keywords:** Biopharmaceuticals, fluorescence, stability, red edge excitation shift, monoclonal antibody

## Abstract

Among the major challenges in the development of biopharmaceuticals are structural heterogeneity and aggregation. The development of a successful therapeutic monoclonal antibody (mAb) requires both a highly active and also stable molecule. Whilst a range of experimental (biophysical) approaches exist to track changes in stability of proteins, routine prediction of stability remains challenging. The fluorescence red edge excitation shift (REES) phenomenon is sensitive to a range of changes in protein structure. Based on recent work, we have found that quantifying the REES effect is extremely sensitive to changes in protein conformational state and dynamics. Given the extreme sensitivity, potentially this tool could provide a ‘fingerprint’ of the structure and stability of a protein. Such a tool would be useful in the discovery and development of biopharamceuticals and so we have explored our hypothesis with a panel of therapeutic mAbs. We demonstrate that the quantified REES data show remarkable sensitivity, being able to discern between structurally identical antibodies and showing sensitivity to unfolding and aggregation. The approach works across a broad concentration range (*μ*g–mg/ml) and is highly consistent. We show that the approach can be applied alongside traditional characterisation testing within the context of a forced degradation study (FDS). Most importantly, we demonstrate the approach is able to predict the stability of mAbs both in the short (hours), medium (days) and long-term (months). The quantified REES data will find immediate use in the biopharmaceutical industry in quality assurance, formulation and development. The approach benefits from low technical complexity, is rapid and uses instrumentation which exists in most biochemistry laboratories without modification.

## Introduction

Maintenance of function and hence efficacy is an important consideration for the developability of biomolecules. This is driven in part by the retention of a native dynamic profile (native flexibility and dynamics) of the protein.^1^ For example, perturbation of enzyme dynamics affects the activity of a large number of enzymes^2–5^ and protein flexibility and dynamics are being exploited for drug design^6^ and protein engineering.^7,8^ A key example of the biological importance of a protein’s dynamic profile lies in antibody epitope recognition. The affinity of antibodies for an epitope is intimately linked to the native protein dynamics.^9,10^ There is also evidence that a protein’s stability is linked to its dynamic profile.^11^ However, the normal dynamic profile of biomolecules is extremely labile and it is very common for antibodies to become inactive or to aggregate, for example on minor temperature variation. This issue is a key concern for the development of biopharmaceuticals, which represent a multi-billion dollar market.^12^

The challenges of developing both stable biomolecule formulations and monitoring for retention of conformation is crucial to the commercial viability and efficacy of biopharmaceuticals. However, capturing subtle or even major changes to a proteins native dynamic profile is challenging. Potential approaches that capture this information include NMR,^13^ EPR,^14^ single molecule spectroscopy,^15,16^ ion mobility mass spectrometry (IM-MS)^17^ and hydrogen/deuterium (H/D) exchange mass spectrometry.^18^ However, at present these approaches are not in routine use due to significant technical complexity and feasibility, instrument expense, time of assay, complex sample preparation and need for specialist analysis. Instead, a breadth of lower resolution approaches such as far-UV circular dichroism (CD), used to detect changes in protein secondary structure, and light scattering or size exclusion chromatography (SEC), are used to detect aggregation. These lower resolution approaches have the advantage that the time to result is much more rapid and requires less technical complexity, but because the information content is lower, one needs to apply a large number of such techniques to gather a full picture.

A protein’s dynamic profile is defined by a free energy landscape (FEL).^19,20^ The FEL can be thought of as a series of energetic hills and valleys that defines the energy required for a protein to fold and change conformation. A well folded, stable protein will occupy an energetic minimum on the FEL, meaning a relatively large amount of energy is required to unfold the protein. In contrast, proteins that are shifted up the energy scale on the FEL tend to be more flexible and dynamic because they are able to sample a range of energetic minima, reflecting different conformational sub-states. However, a consequence is that they may be less thermodynamically stable. Any protein conformational change, by definition, must therefore be accompanied by either a change in the protein FEL or a transition to new minima on the FEL.

Tryptophan (Trp) residues in proteins are extremely sensitive reporters of the immediate molecular environment.^21–23^ Trp residues can display a shift in their emission maximum with a decreasing energy of excitation, because the lower energy photons selectively excite discrete conformational states of the Trp-solvent system, the so called red edge excitation shift (REES) effect.^24–28^ The REES effect has primarily been used to distinguish between folded and unfolded states of proteins.^29–33^ We have demonstrated that by quantifying the REES data more directly (described below), the Trp REES effect (Figure 1A) becomes a powerful tool that informs on the dynamic profile of a protein.^34,35^ Specifically, the quantified REES data reflect the equilibrium of protein conformational states characterised by a proteins free energy landscape (FEL).^34^ More recently we have shown a similar detection sensitivity in multi-trp proteins, with the approach being able to discern differences in molecular flexibility, where the X-ray crystal structures are identical.^35^ Similarly, we have recently shown that extrinsic fluorophore labelling or mAbs can be used to track changes in low-*n* oligomer formation.^36^ We term our method of quantifying REES data, QUBES (quantitative understanding of bimolecular edge shift), in order to distinguish other treatments of REES data. Based on the apparent power and potential information content of the REES phenomenon, we reason that this approach could be developed further to deliver sensitive detection of changes in a proteins dynamic profile as well as overall conformation. Such a tool would then have utility in the discovery and development of mAbs.

**Figure 1.**
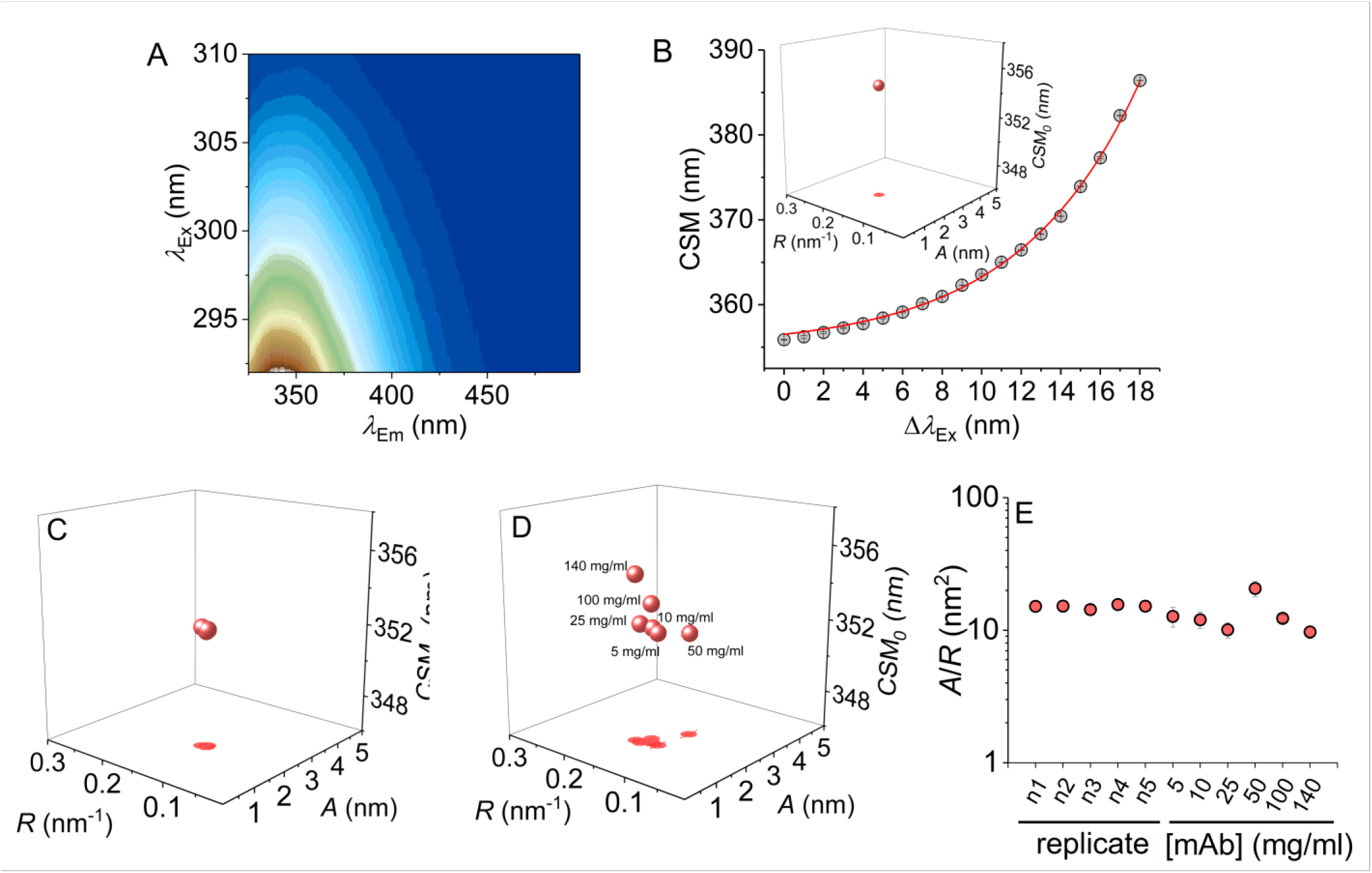
QUBES data using Eq 2 and consistency of datasets. **A,** The parameters in Eq 2 are extracted from a combined excitation-emission spectrum for protein Trp residues. **B,** The CSM *versus* excitation wavelength (grey data) is fit to Eq 2(solid red line), to give a single data point governed by 3 parameters (shown in *inset*). **C,** QUBES data for five biological replicates of mAb1, each having three technical replicates. **D,** The effect of variation in mAb 1 concentration. **E,** Comparison of *A*/*R* values extracted from panels C and D. *Conditions*, Histidine pH 5.6, 15 °C, buffer, 5 mg ml^−1^ (panel C).

Herein we demonstrate that quantifying the REES effect could be used as a simple spectroscopic fingerprint to assess the stability and structure of mAbs. We rigorously test our hypothesis using both commercially available therapeutic mAbs and examples from those in active commercial development. Our findings suggest the approach could have value as part of quality control of existing therapeutic mAbs and enhancing decision making during mAb development.

## Methods and Methods

### REES data collection and analysis

All fluorescence measurements were performed using a Perkin Elmer LS50B Luminescence Spectrometer (Perkin Elmer, Waltham, MA, USA) connected to a circulating water bath for temperature regulation (± 1 °C). Samples were incubated for 5 minutes at the given conditions prior to recording measurements. Measurements were performed at 10°C, unless otherwise stated. Excitation and emission slit widths were 5 nm. Tryptophan emission was monitored from 325 to 500 nm. The excitation wavelength was subsequently increased in 1nm steps for a total of 19 scans. 3 sets of individual scans were averaged. The corresponding buffer control was subtracted from the spectra for each experimental condition and this also removes the Raman peak water peak. The center of spectral mass (CSM) was calculated using the following equation:

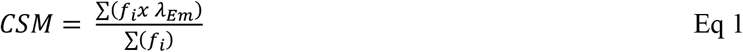

Where *f*_i_ is the measured fluorescence intensity and *λ*_em_ is the emission wavelength. We would stress the importance of using a consistent wavelength range when reporting CSM data, as the magnitude will be dependent on the wavelength range chosen. The data are extracted by fitting the CSM *versus λ*_Ex_ data as described in the manuscript. Data fitting and plotting was performed using OriginPro 2016 (Microcal).

### Antibody samples, unfolding and aggregation

Therapeutic antibodies (Figure 2A) were provided by Bath ASU and were either extensively dialysed (for urea denaturation experiments) or diluted into Tris-Cl buffered saline pH 8. All buffer components were of a spectroscopic grade. Antibody denaturation was achieved by extensive dialysis into a buffered solution of 8M urea or 0M urea as a control. Antibody aggregation was achieved through incubation at elevated temperatures and monitored by DLS as described in the manuscript.

**Figure 2.**
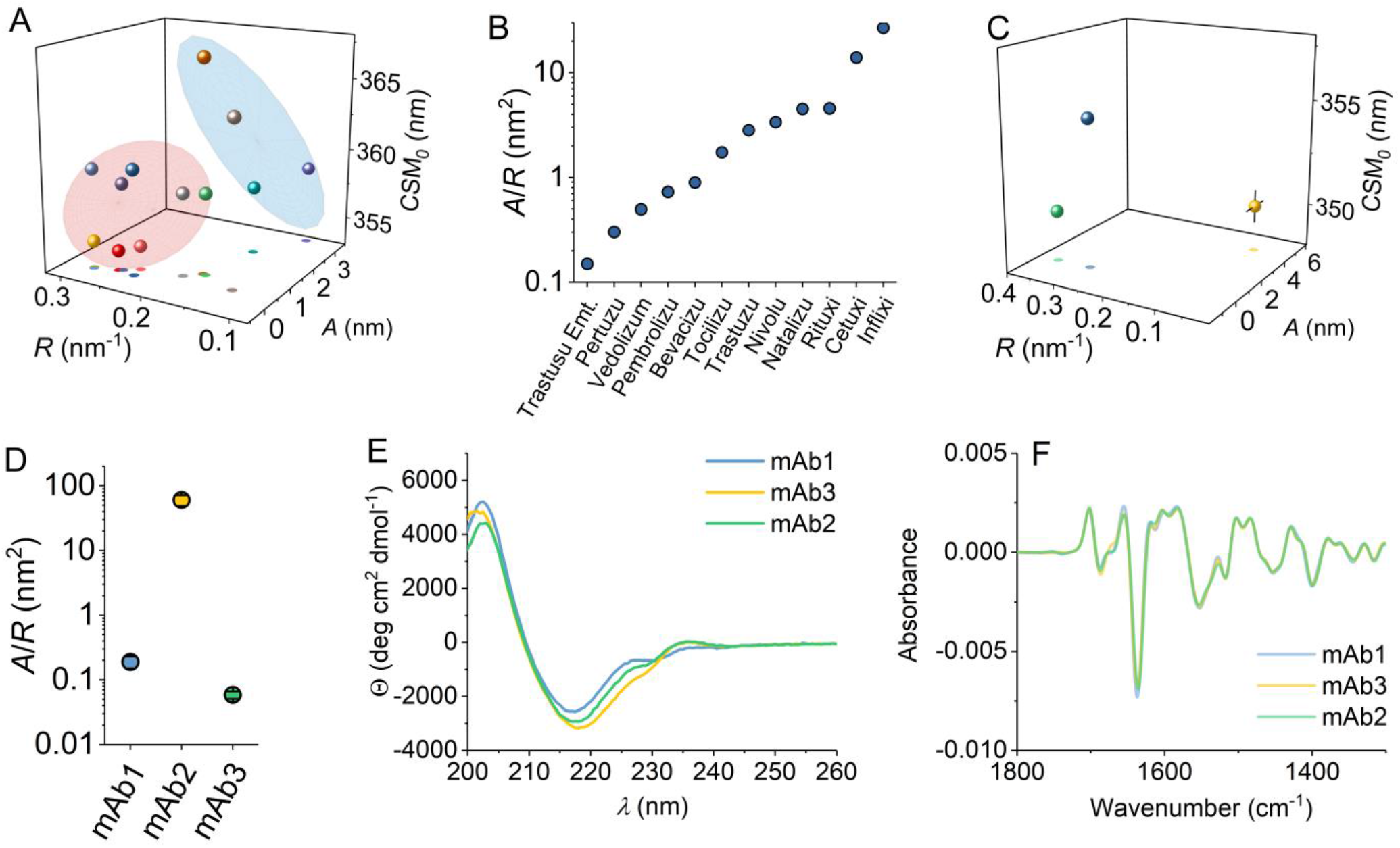
**A,** QUBES data for a series of zumab (red contour), ximab (blue contour) and lumab (grey) examples. Coloration as Pembrolizumab (green), Vedolizumab (blue), Pertuzumab (orange), Natalizumab (yellow), Bevacizumab (Indigo), Trastuzumab (red), Trastuzumab emtansine (light blue) and Tocilizumab (light green), Nivolumab (grey), Rituximab (gold), Inflximab (purple) and Cetuximab (emerald). **B,** Comparison of *A*/*R* values extracted from panel A, ranked from smallest to largest value. **C,** QUBES data for mAbs in commercial development; mAbs1-3, shown as blue, yellow and green, respectively. **D,** Comparison of *A*/*R* values extracted from panel C. **E,** CD data for mAbs1-3. **F,** and FTIR (second derivative) mAbs1-3. *Conditions*, for the mAbs in panel A; 50mM Tris pH 8, ~ 1mg/ml at 10°C. For the mAbs in panel C; Histidine pH 5.6, 15 °C, buffer, 5 mg ml^−1^ (panel C).

### Structure-based calculations

Partial Fab region structures (X-ray crystal structures) were used for all structure-based calculations. To ensure comparability and the presence of HVL regions we homology modelled each structure using the RosettaAntibody Online Server. The model with the lowest calculated energy was selected for further analysis. The solvent accessible surface area (SASA) for the tryptophans in each mAb was calculated using Pymol (Delano Scientific). Normal mode analysis (NMA) was undertaken for each mAb using the elNémo online server. The frequency of each non-trivial normal mode was recorded for modes 7-106.

### Rigidity analysis

Pebble-game rigidity analysis^37^ divides a protein structure into a number of rigid clusters (RCs) depending on the distribution of constraints in the system. The results depend particularly on the inclusion of hydrogen bonds in the constraint network, which is controlled by an energy cutoff parameter *E*_cut_. As *E*_cut_ is decreased from zero to negative values, weaker hydrogen bonds are excluded from the constraint network and the structure becomes less rigid. We track the rigidity by considering the fraction (*F*) of main chain residues which lie within the *N* largest rigid clusters (for *N* = 1 to 20) for *E*_cut_ values lying in the range from 0 kcal mol^−1^, where the structure is largely rigid, to −4 kcal mol^−1^, where the structure is largely flexible.^38^ In order to obtain a single parameter describing the rigidity, which can be used to compare the different antibody structures in our study, we average the rigid fraction F over both N and *E*_cut_. We term this overall value the sum value of rigid clusters, SVRC. The higher the value of SVRC, the more rigid the structure.

## Results and Discussion

### Quantifying the REES effect for multi-Trp proteins gives a unique ‘fingerprint’ for different mAbs

We have monitored the edge-shift effect for a range of mAbs, shown in Figures 1 and 2. The data is collected from the combined excitation and emission spectrum, monitored for each mAb, giving a high information content fluorescence data set (example shown in Figure 1A). The intensity and peak position of the emission incorporates information on (i) the number of Trp residues in the sample (ii) the degree of solvent exposure of the Trp residues^21^ including arising from different rotamers (iii) energy transfer to the peptide backbone,^22^ (iv) homotransfer to other Trp residues^23^ and (v) photoselection of discrete solvation environments at low energy excitation.^24,25^ We have previously shown that contributions from Tyr emission are essentially negligible over the excitation range used.^34^ From these data (Figure 1A) one can extract the variation in the emission spectra with excitation wavelength as either the change in the emission peak position (*λ*_max_) or as the change in the centre of spectral mass (CSM). We prefer the use of CSM as it does not require model fitting to accurately extract the emission peak maximum and incorporates information on the whole data-set. Figure 1B shows an example of the resulting plot of CSM *versus* excitation wavelength for an example mAb.

The data in Figure 1B show a curved relationship and this is typical and similar to reports with proteins containing single Trp residues (for example ref 34).^34^ Simple linear fitting of these data is clearly inadequate to capture the full information content contained in the data set. We have previously fit an exponential function to these data to capture the information contained in the curvature of the edge-shift data,^35^

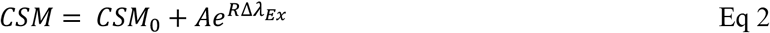

Where CSM_0_ is the CSM value independent of the excitation wavelength, *λ*_Ex_, determined by the amplitude, *A*, of an exponential with a curvature determined by *R.* The plot of the resulting parameters yields a single three-dimensional data point (Figure 1B, *inset*), which is a direct quantification of the extremely complex spectral fingerprint shown in Figure 1A. We term these data, QUBES (quantitative understanding of bimolecular edge shift) data, to distinguish from other analyses. We have previously found that the ratio; *A*/*R* from Eq 2, gives a single value that appears to relate to changes in protein flexibility^35^ and is a simple visual metric of changes in the QUBES data. Specifically, large *A*/*R* values reflect more flexible proteins and small *A*/*R* reflect a more rigid protein (as defined above). Note that the CSM_0_ value is also key to data interpretation and we consider the use and interpretation of the data in detail below

We have established the reproducibility of the QUBES data, using an example IgG4 mAb (mAb1) that is in active development and the resulting data for five biological replicates is shown in Figure 1C. The data show extremely small variance and are essentially the same within the extracted error values (Eq 2). Specifically, the average values and standard deviations of the parameters extracted from Eq 2 are *A* = 2.0 ± 0.03, *R* = 0.13 ± 0.003 and *CSM*_0_ = 352.3 ± 0.07. We have also explored the variation in the data with respect to a large range of protein concentrations (5 mg/ml – 140 mg/ml), Shown in Figure 1D. At ‘low’ concentrations (5-25 mg/ml) we find relatively little variance in the REES data quantified using Eq 2 (Figure 1E). However, as the concentration increases there are shifts in the parameters, particularly at very high concentrations (100 – 140 mg/ml), primarily manifesting an increase in CSM_0_ and to a lesser extent with the other parameters (Figure 1E). Specifically, the average values and standard deviations of the parameters extracted from Eq 2 for the data in Figure 1D are *A* = 1.84 ± 0.4, *R* = 0.15 ± 0.01 and *CSM*_0_ = 352.7 ± 1.2. It is important to note that we have used a flash-lamp-based fluorimeter and so the power is relatively low on excitation. That is, we do not observe any appreciable photo-bleaching across triplicate measurements of the same mAb and this is the case for all mAbs used in this study.

Given the consistency of the values within replicates (Figure 1C), potentially the variance in the parameters with respect to concentration may reflect changes in structure/stability. For example, increasing protein concentration will increase the propensity for aggregation, but at higher concentrations one expects macromolecular crowding and viscosity variance to affect protein flexibility.^34,35^ We note that at elevated protein concentration, the inner filter effect, will be present. However, the REES effect is independent of the magnitude of fluorescence emission and only relies on changes in the structure of emission spectra. Below we develop the understanding of the information content of the data with respect to mAb structure and stability.

We have measured QUBES data for a range of commercially available therapeutic mAbs. These mAbs represent different classes including, chimeric (ximab; Rituximab, Infliximab and Cetuximab), humanised (zumab; Pembrolizumab, Vedolizumab, Pertuzumab, Natalizumab, Bevacizumab, Trastuzumab, Trastuzumab emtansine and Tocilizumab) and human (lumab; Nivolumab) in the same buffer system, shown in Figure 2A. That is, each mAb is not in it’s commercial formulation so that more valid comparisons can be made. From Figure 2B, we find that there is a difference in the extracted values (using Eq 2) for each of the mAbs studied and between classes of mAb. Similar to the example given in Figure 1C, the individual values from each of the parameters in Eq 2 are extremely reproducible both for individual replicates of the same sample and also batch-to-batch variation, with a typical standard deviation. As such, the differences we monitor in Figure 2A are *bone fide* and do not represent the absolute variance across the samples as a whole.

The separation of the QUBES data (Figure 2A and 2B) is interesting given the very high sequence conservation of the mAbs and the overall structural similarity. Nine of the twelve examples we have studied are of the IgG1 isotype except Pembrolizumab, Vedolizumab and Natalizumab, which are IgG4, differing only by three residues in the hinge region and retaining the same inter-heavy chain disulphide bonds. We reason that for the same class of mAb (chimeric, humanised or human) the three-dimensional structures can be considered to be essentially identical. For example, the far-UV circular dichroism spectrum and dynamic light scattering (DLS) profile of these full length mAbs is highly similar if not identical (see below) as expected for proteins with high sequence similarity and similar overall structures particularly in the percentage of secondary structure content. Indeed, we have previously found that REES data can vary significantly in a multi-Trp protein for a single amino acid variant with identical overall structure.^35^

To make this point explicit, Figure 2D-G shows the comparison of QUBES data (Figures 2D and 2E) and the corresponding far-UV CD and FTIR (Figures 2F and 2G) spectra for three IgG4’s (mAb1-3) that are in development. As with the commercially available examples given in Figure 2A, we see significant differences in the QUBES data for mAb 1-3 (Figures 2D and 2E). However, the structures are essentially the same as assessed by the essentially invariant CD and FTIR data (Figure 2F and 2G).

### The QUBES data are sensitive to mAb flexibility

The data above prompt the question; why is the QUBES data different for essentially identical protein structures? Differences in the number and position of Trp residues might give rise to different QUBES data. The Fab region [*Supporting* information (SI), Figure S1] of the mAbs, shown in Figure 3A, contains the most sequence variability and there are some small differences in the number and/or position of Trp residues for some of the mAbs in the Fab region (Figure S1 and S1B and Table S1). However, these differences (number of residues and solvent accessible surface area) do not show an obvious correlation with the extracted values from Eq 2 (Figure S1C and S1D). These small differences would not therefore appear to be sufficient to explain the differences in the data shown in Figure 2B.

**Figure 3.**
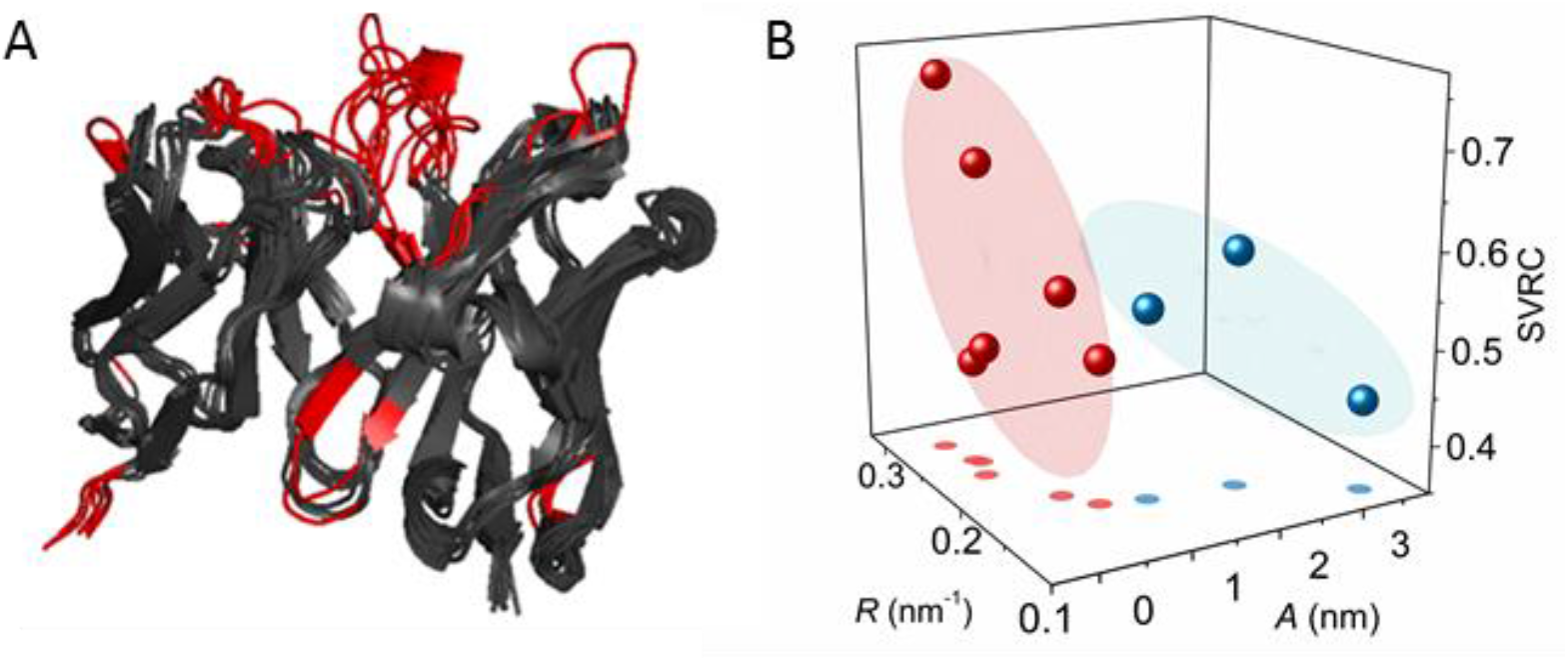
QUBES data reflects changes in molecular flexibility. **A**, Structural overlay of the heavy and light chains of the Fab for the zu- and ximabs. Regions shown in red show significant variation from the overlaid structures. **B**, relationship between the values extracted from Eq 2 (parameters *A* and *R*; red for zumab, blue for ximab and the calculated molecular flexibility (sum value of rigid clusters; SVRC). The coloured planes represent the 30% confidence interval for the data sets and are to aid the eye.

We have previously shown that the curvature in the REES effect captures information on the equilibrium of conformational states accessible on the FEL, which can be thought of as the proteins relative rigidity/flexibility.^34,35^ This level of discrimination is achieved even for multi-Trp proteins as along as the structure and number of Trp residues is the same/similar. Despite the overall structural similarity of the mAbs, we reasoned that the mAbs may have significantly different flexibilities at a range of time and length scales; ranging from natural ‘breathing’ motions of the whole protein to more rapid conformational sampling by hypervariable loop (HVL) regions of the Fab (Figure S1).

To explore the potential correlation between the QUBES data and mAb flexibility, we have turned to computational calculations, using pebble-game rigidity analysis^37^ to assess the structural rigidity of the partial Fab regions. This approach provides information on the relative stability and rigidity of comparable protein structures as we have described previously.^35,38,39^ From this analysis we extract a parameter describing the overall rigidity of the protein^37^ (sum value of rigid clusters; SVRC, see *Methods*) as shown in Figure S2. As a generalisation, the larger this value, the more rigid the protein.

Figures 3B and 3C show the correlation between the values extracted from fitting to Eq 2 and the extracted rigidity for those mAbs with available high-resolution crystal structures. We have separated the values between the humanised and chimeric antibodies. From Figure 3B, the SVRC values vary very significantly, suggesting major differences in rigidity despite high structural similarity. Moreover, we find a clear correlation between the *A* and *R* values with the calculated rigidity (SVRC) for both chimeric and humanised antibodies (Figure 3C). This correlation suggests that a small *R* value and a large *A* value (increased *A*/*R* ratio) are indicative of increasing molecular flexibility, i.e. less rigid structures, at least for the mAbs studied here. These data suggest the reason for the separation of the extracted values arises in large part from the difference in molecular flexibility of the mAbs and this is consistent with our previous findings for both single and multi-Trp proteins.^34,35^

### Probing the information content and sensitivity of QUBES data

Changes in protein structure are accompanied by a change in the equilibrium of conformational states. We have recently demonstrated that monitoring REES for mAbs labelled with an extrinsic fluorophore can report on relatively low-*n* changes in oligomeric state.^36^ Similarly, the REES effect has been used to infer differently folded/aggregated states of proteins using singe-Trp containing examples.^30^ Based on these findings and given the sensitive discrimination of mAbs by the QUBES data shown above, we wished to explore whether differences in multi-Trp REES data are sensitive to changes in protein structure in mAbs. We have therefore tested whether the QUBES data can be used to identify unfolded and/or aggregated states of the mAbs. The mAbs were subjected to stress conditions that would promote unfolding (8M urea) and aggregation (65 °C ~5 hrs). Note that we have used a buffer with elevated pH (pH 8) compared to the therapeutic formulation, to increase the rate of aggregate formation. Otherwise the process would take ~days-weeks and be difficult to monitor and control.

Figure 4A shows the data extracted from fitting REES data to Eq 2 for these conditions and Figure 4B shows the data simplified to show just the *A*/*R* value as above. Incubation with urea will cause the mAbs to partially unfold (retaining the native disulphide linkages), but also prevent aggregate formation, whereas thermal denaturation, particularly for the mAbs studied, directly drives aggregate formation. We monitor the formation of soluble aggregates in our thermally denatured mAbs by dynamic light scattering (DLS), shown in Figure S3.

**Figure 4.**
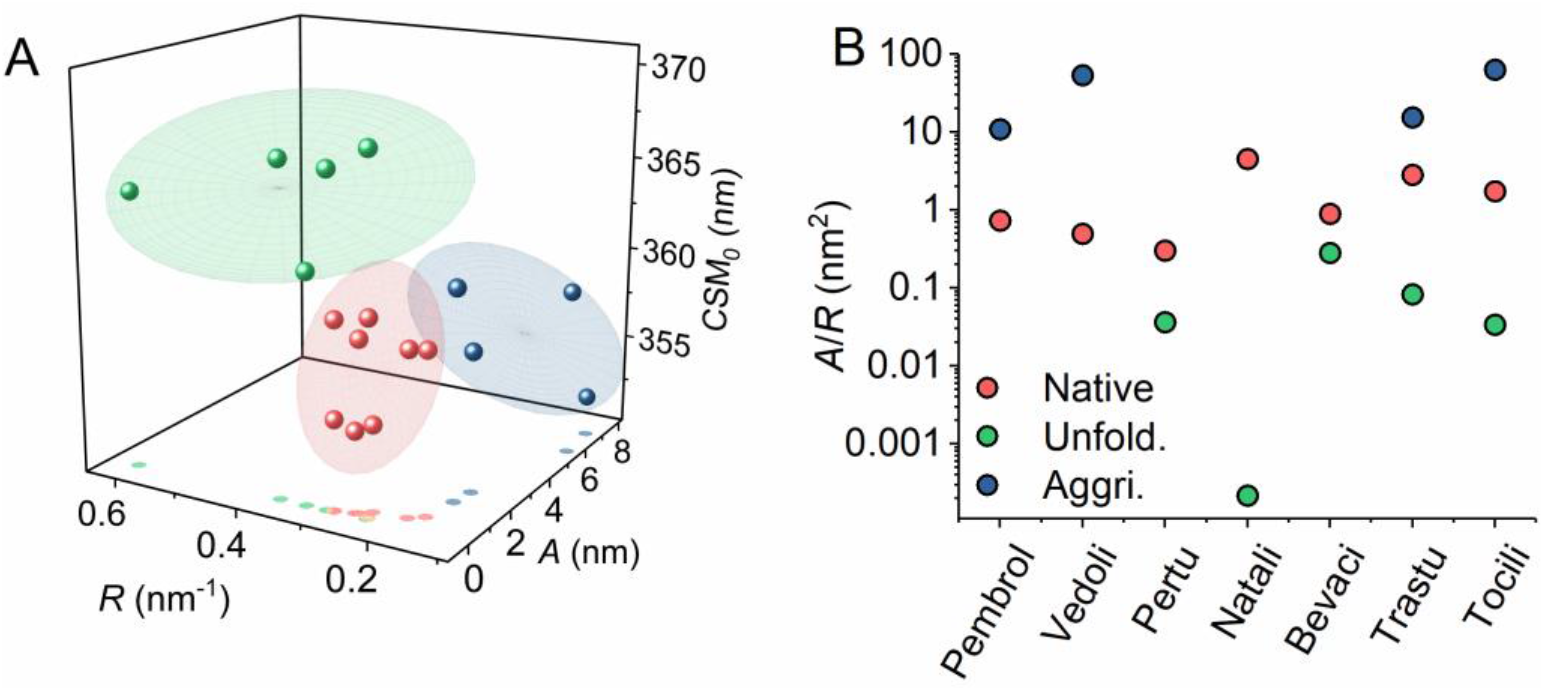
**A,** The QUBES data can be used to accurately reflect and differentiate between mAb unfolding and early stage aggregate formation. Extracted values for zumab shown in Figure 2B (red), incubated in 8M urea (green) and thermally aggregated (purple). **B,** Resulting *A*/*R* values from Panel A.

Figure 4A and 4B shows that the unfolded (urea denatured) mAbs cluster to higher CSM_0_ and *R* values, but smaller *A* values. These data reflect a flatter REES effect of a smaller total magnitude based on a simple linear fit. Based on our findings from model protein studies of the REES effect,^20^ we would suggest these data reflect more solvent exposed Trp residues (indicated by the higher CSM_0_) and a decrease in the range of conformational states available to the protein as it tends towards a single state; a completely unfolded linear polypeptide chain. These notions are in-line with the range of proteins that have been observed to have a decreased REES effect upon unfolding, which reflects the transition towards a restricted equilibrium of conformational states characterised by a fully unfolded protein. ^24–26^ Conversely, treating the gross trends in the whole data set (Figure 4A and 4B), we find that thermally denatured mAbs cluster to elevated *A* values, but smaller *R* and CSM_0_ values (we explore these changes in more depth below). Based on the corresponding DLS data (Figure S3), this shift in the QUBES data would therefore seem to predominately reflect the formation of soluble aggregates.

We therefore find that the REES data quantified using Eq 2 is not only able to discern native and denatured protein but also to separate proteins that are unfolded from those that are aggregated. We suggest that the observed variance in the extracted values for the denatured mAbs (Figure 4A) may reflect the differing extent of unfolding or aggregation for each of the samples and the specific nature of the unfolded or aggregated states.

To explore the sensitivity of the QUBES data we have studied the response to a large range of physical and chemical perturbations that are commonly experienced by mAbs during a manufacturing process. To test the robustness of therapeutic mAbs for developability, they are subjected to forced degradation studies (FDS) with stress conditions including temperature variation, effect of chemical modification (deamidation/ oxidation), agitation, light exposure and pH variation. Figure 5A shows the QUBES data for a range of stresses applied to mAb1 over 14 days. Figure 5B shows the corresponding *A*/*R* values ranked for the notional decrease in stability (low to high values as above). From figure 5A and 5B, the QUBES data shows sensitivity to the full range of different physical and chemical stresses. From Figure 5B, the three physical perturbations that have the largest effect on mAb 1 stability are high temperature (50 °C), low pH (pH 3) and high light exposure (5 mlux.hr). The shifts to larger *A*/*R* ratios in the stressed samples are indicative of increased flexibility, while the significant increase in CSM_0_ observed for the 50°C, pH 3 and 5 mlux.hr samples suggests an increase in the average solvent exposure. The detection of unfolding in the pH 3 sample is not surprising, as there is evidence that low pH can cause unfolding of the Fc for IgG4s^(40)^ but see our discussion below.

**Figure 5.**
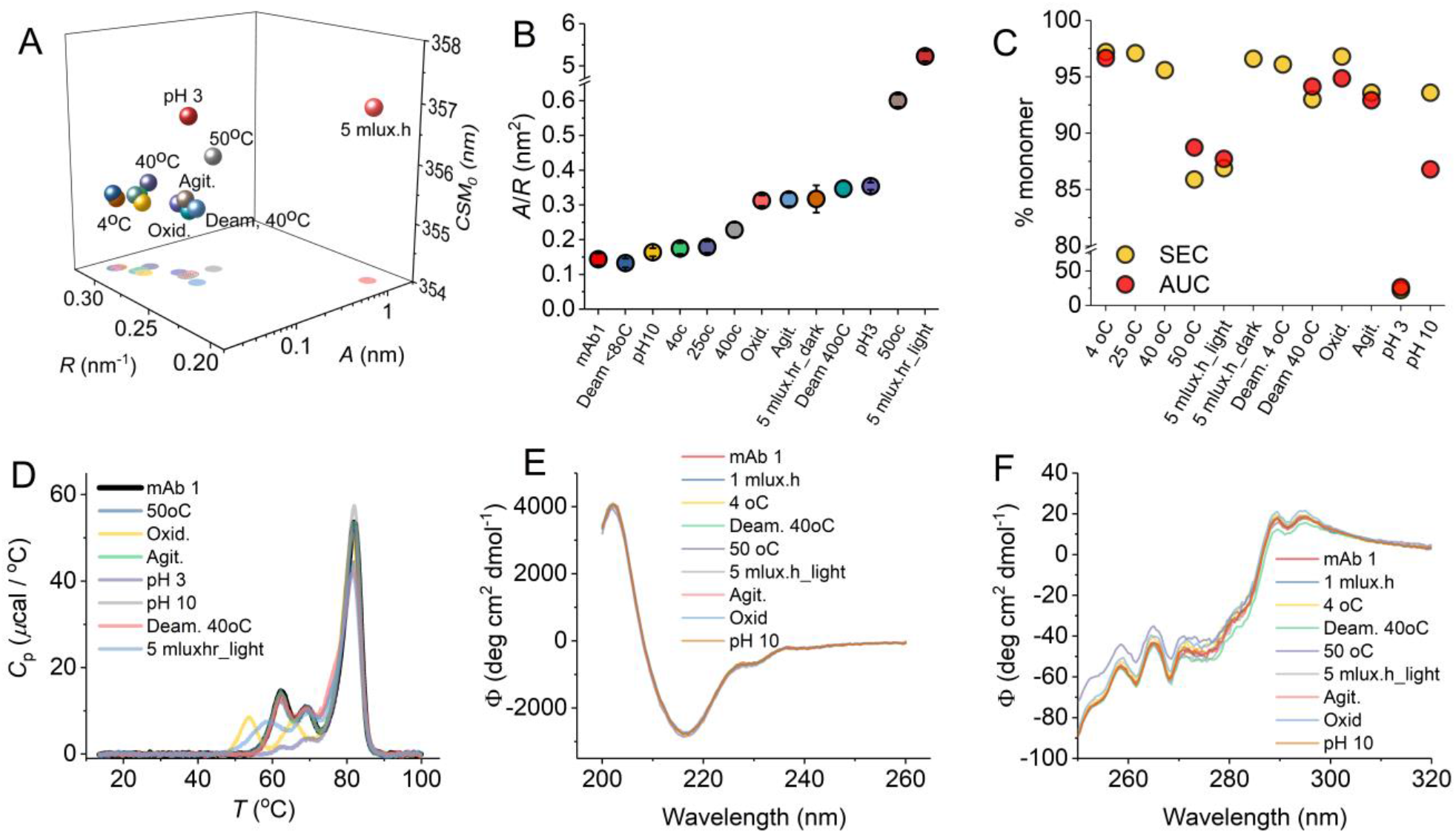
Sensitivity of QUBES data to different physical perturbations. **A,** QUBES data for mAb 1 subjected to a range of physical perturbations as part of a forced degradation study. **B,** Comparison of *A*/*R* values extracted from panel A. **C,** Monitoring changes in aggregation using SEC (yellow), CGE (blue) and AUC (red). AUC monomer % is inferred from the fraction of aggregated material detected. Grey data are the average of the three techniques and the error bars are the standard deviation. **D,** DSC data for different FDS perturbations. **E,** Far-UV CD spectra for different FDS perturbations. **F,** Near-UV spectra for different FDS perturbations. *Conditions,* Samples were incubated at the indicated conditions for two weeks. QUBES measurements were performed at 15 °C in formulation buffer, at 5mg ml^−1^.

Figure 5C-5F show a range of other approaches used to assess the effect of different physical perturbations on mAb1. These include assessment of aggregation [Figure 5C; size exclusion chromatography (SEC) and analytical ultracentrifugation (AUC)], stability to unfolding [Figure 5D; differential scanning calorimetry (DSC)] as well as secondary and tertiary structure content [Figures 5E and 5F; far-UV and near-UV circular dichroism (CD), respectively]. From these data the perturbations that affect mAb1’s aggregation propensity the most (outside of calculated error) are the same as suggested by the QUBES data above. That is, high temperature (50 °C), low pH (pH 3) and high light exposure (5 mlux.hr). Similarly, the DSC data (Figure 5D) suggest the perturbation that affects unfolding thermodynamics the most are low pH (pH 3) and high light exposure (5 mlux.hr), but now oxidation instead of high temperature. Finally, the far-UV CD data show essentially no variance under any condition, suggesting the secondary structures are essentially unaffected within the limitations of the detection of the approach. There are potentially some very subtle differences in the near-UV CD data, particularly for the 50 °C condition, but we are cautious interpreting these data because the differences are small. Note that the pH 3 sample precipitated upon exchange into the buffer used for CD analysis and so no data were collected for this sample.

We have posited (above) that the approach reflects changes in the available equilibrium of conformational states; the molecular flexibility. For the FDS studies above, the QUBES data will additionally be convolved of heterogeneity arising from different aggregate conformations as well as changes in molecular flexibility of the remaining monomer population. That is, the contributions to the QUBES data will be complex. However, From Figure 5, we find that not only are the QUBES data sensitive to a range of physical perturbations, they give complementary results to a breadth of other techniques that are commonly used.

### QUBES predicts protein stability

Our data suggest that the QUBES data are remarkably sensitive and that by capturing information regarding changes in protein flexibility the approach could have excellent utility when combined with complementary approaches. We hypothesise that the quantification of the curvature in REES data might reflect information on protein flexibility (see below). Increased protein flexibility is typically correlated with decreased thermodynamic stability because there is a smaller energetic barrier of unfolding as evidenced from a range of mesophile *versus* thermophile enzyme studies^41^ and as we have demonstrated recently.^35^ There is also evidence from HDX-MS that flexibility may be linked to long term stability of mAbs.^11^ We therefore now ask whether, given the nature of the putative detection sensitivity of the QUBES data, can we use the approach to predict changes in protein stability?

Figure 2B shows the commercially available mAbs ranked based on the calculated *A*/*R* value. These data are notionally a simple visual metric of differences in flexibility. If so, we would expect that the *A*/*R* value would track with the stability of a protein and so could potentially be predictive of changes in stability. To test the potential for the QUBES data to be used in a predictive manner we have therefore explored the thermal stability of three commercial mAbs with *A*/*R* values that suggest increasing flexibility and therefore an inferred decrease in stability.

The QUBES data for Pertuzumab, Vedolizumab and Nivolumab are shown in Figure 6A. From Figure 2C, Pertuzumab is predicted to be the most stable and Nivolumab the least stable as assessed by the relative increase in the *A*/*R* value (Figure 6B). We stress this trend is for these mAbs in the same buffer system, not their commercial formulations. We incubated these mAbs at 65 °C for 5 hours in a pH 8 buffer and observed the formation of aggregates via dynamic light scattering (DLS). We opted for DLS to immediately capture the aggregation state and so we could directly correlate with the QUBES data. In all cases, heating induced an increase in the *A*/*R* value suggesting a transition to a less stable protein. Similarly, from Figure 6A we see a decrease in the *CSM*_0_ parameter as with our previous findings for aggregated mAbs (Figure 4).

**Figure 6.**
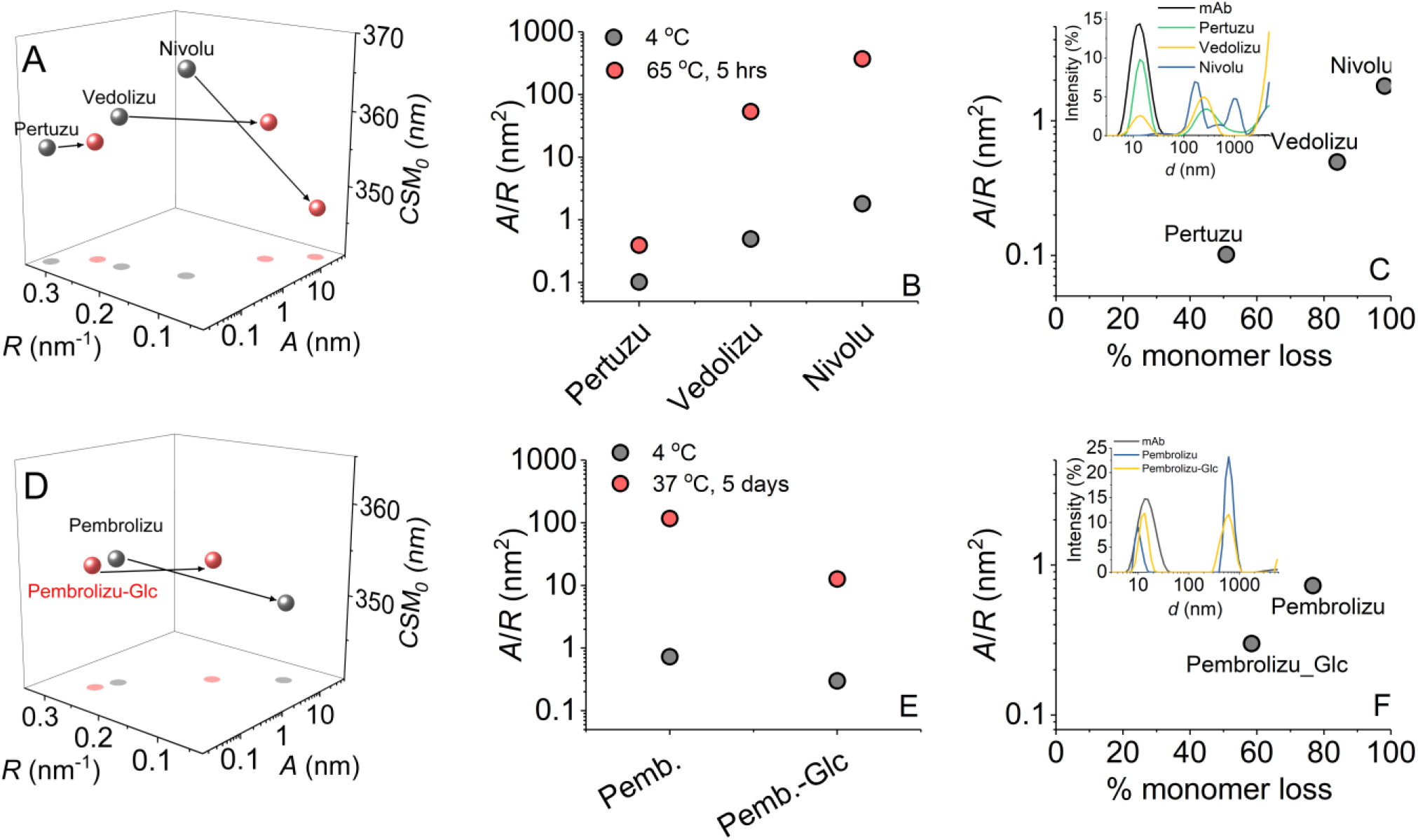
QUBES data can predict thermodynamic stability. **A,** Extracted QUBES data values for three therapeutic mAbs predicted to have different thermodynamic stabilities. Grey data are for the native protein, red data are after incubation at 65 °C for 5 hours. **B,** Comparison of *A*/*R* values extracted from panel A. **C,** Comparison of *A*/*R* value from panel B to the fractional loss of monomer as assesed by DLS (inset). **D,** Extracted QUBES data values for Pembrolizumab with and without glucose present Grey data are for the native protein, red data are after incubation at 37 °C for 5 days. **E,** Comparison of *A*/*R* values extracted from panel D. **F,** Comparison of *A*/*R* value from panel E to the fractional loss of monomer as assesed by DLS (inset). *Conditions,* 50mM Tris pH 8, ~ 1mg/ml at 10°C

The resulting correlation between the fractional loss of monomer (assessed by DLS; Figure 6C inset) and the *A*/*R* value is shown in Figure 6C. Based on our QUBES data (Figure 6A and 6B) and the corresponding DLS profiles (Figure 6C) we find that there is a trend for a more significant fraction of soluble aggregate present for Nivolumab compared to Vedolizumab compared to Pertuzumab. These data therefore confirm our hypothesis that the extracted values from Eq 2 can be used in predictive manner to infer the relative thermodynamic stability of a sample. We note that the predictive power is only appropriate for the same sample, since each protein will exhibit a specific spectral ‘fingerprint’ (Figure 1A) signature. In the present case, the mAbs have three dimensional structures that are essentially identical and so the comparison between them is valid.

We next wished to explore the potential of the QUBES data for formulation of stable biopharmaceutical preparations and stability over longer timescales (days). To that end we have monitored the temperature induced unfolding and aggregation of Pembrolizumab both in the presence and absence of a known adjuvant (glucose). The resulting QUBES values and DLS profiles are shown in Figure 6D and 6F, respectively. Based on the shift in the *A*/*R* value on addition of glucose (Figure 6E) we would predict that the glucose should have a stabilising effect on Pembrolizumab. From Figure 6F, we find that incubation of Pembrolizumab at 37 °C for 5 days induces significant formation of soluble aggregates. However, as we predict, glucose provides significant protection from aggregate formation with a lower percentage of soluble aggregate formation as assessed by both the QUBES data and DLS profiles (Figure 6E). We note that we do not observe any post-translational modification of the mAbs (glycation) based on a fluorescence reporter system^42^ and so the effect is due to stabilisation of the mAbs and not an artefact arising from glycation.

Given the QUBES data appears to show predictive capacity for stability on the hours-days timescale, we wished to assess if the approach could show sensitivity to stability over much longer term storage conditions. To that end we have monitored the fractional loss of monomer for mAbs 1-3 (Figure 2D) using SEC. From Figure 2E, the *A*/*R* value suggests that mAb 3 will be the most stable and mAb 2 the least stable. Figure 7 shows the correlation between the fractional loss of monomer over 6 months, at 25 °C. From Figure 7B, we find that the QUBES data tracks with the loss of the monomer (as with the examples shown in Figure 6), demonstrating that the QUBES data are able to show remarkable predictive ability, even to the level of fractions of 1 % change in monomer concentration on long-term storage. We would stress that the data we have at present suggests the quantified REES data can reflect relative changes in stability for a specific protein, not define absolute timescales. Potentially this could be achieved on a case by case basis by forming calibration curve for a specific protein.

**Figure 7.**
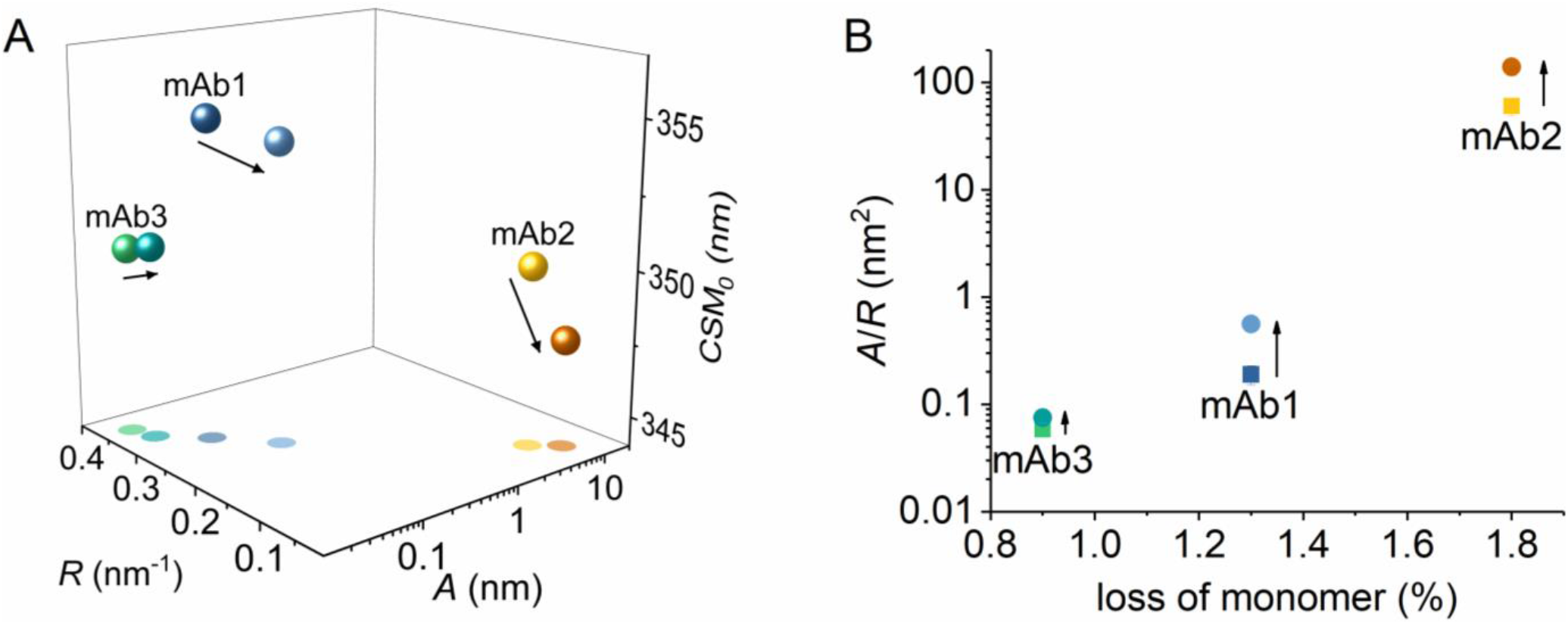
**A,** QUBES data for mAbs1-3 at 15 and 40 °C. Solid arrow indicates the change in extracted values from 15 and 40 °C. **B,** Relationship between the *A*/*R* value and the loss of monomer over 6 months at 25°C. *Conditions,* Histidine pH 5.6, 15 °C, buffer, 5 mg ml^−1^.

Using mAb1-3 we wished to explore the potential of differential temperature measurements as an additional discriminative probe of stability. Increasing the temperature of a protein will alter the distribution of conformational states, providing access to thermodynamic minima on the FEL that would otherwise have a low fractional population. In a simplified sense, proteins should become more flexible as the temperature increases, until the structure begins to breakdown and unfolding occurs, often proceeding to aggregation. Within the temperature range that the protein retains its native folded state, temperature might therefore be a useful discriminating parameter to further characterise the flexibility of a protein by REES. The data in Figure 7 for mAb 1-3, are shown at both 15 and 40 °C. From Figure 7B we find that each mAb shows a distinct *A*/*R* value and these are differently temperature dependent for each of mAb1-3, but in all cases *A*/*R* increases with temperature. These data are an illustration the principle described above, i.e. that increasing temperature should increase protein flexibility and this should manifest as a larger *A*/*R* value.

At least for mAbs 1-3, there does not appear to be a particular temperature that is optimal for detecting stability. That is, the same trend is evident at low and high temperatures. We note that the REES effect itself will be temperature dependent since the dipole moment of the environment will vary with temperature. However, given the similarity in sequence and structure for these mAbs we expect the temperature dependence of the REES effect to be similar and so the comparative data is useful.

## Conclusions

By experimentally monitoring a large number of mAbs we are able to provide a schematic for the interpretation of the QUBES data as shown in Figure 8. Our data above, as well as our previous work^34,35^ suggests that the curvature in the REES effect contains valuable information on the proteins FEL, reflecting differences in rigidity/flexibility. The theoretical basis of the REES effect is predicated on the concept that decreasing energy of excitation can photoselect for discrete species within an equilibrium. The broader this equilibrium the more species can be photoselected and so one anticipates a larger REES effect. In terms of a plot of CSM *versus λ*_Ex_ (e.g. Figure 1B), we expect a bigger absolute magnitude of spectral red shift (reflected in the *A* value from Eq 2) and also more curvature (reflected in *R* value from Eq 2). The reason for this is that a decreasing excitation energy will photoselect for fewer conformational sub-states within the equilibrium.

**Figure 8.**
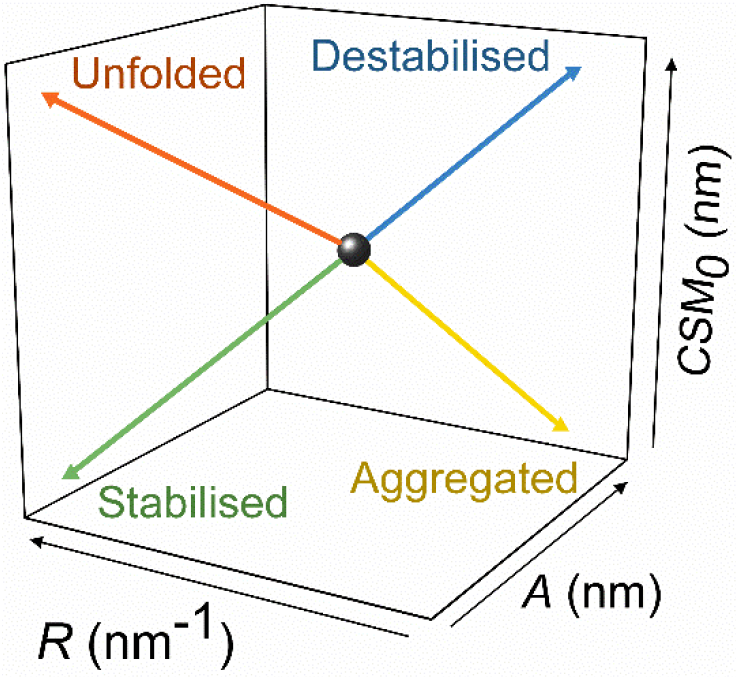
Summary of the detection capability of QUBES data. The change in position of the parameters separatley reflects unfolding and aggregation as well as (de)stabilisation. The stability of the sample is a reflection of the changing molecular flexbility (increasing rigidity providing increasing stabilistation).

We stress that there is potential for an over simplistic interpretation of the data based on the ratio *A*/*R* alone. That is, on urea induced infolding, the *A*/*R* tends to decrease (Figure 4), similar to the putative stabilisation (Figure 6 and 7). Similarly, aggregation tends to increase the *A*/*R* value (Figure 4), as does destabilisation of the protein (Figure 6 and 7). We suggest that the full range of QUBES data should be used to assess a protein, which means including information from the CSM_0_ value. This value represents information on the solvent exposure of Trp residues, with a larger value reflecting a higher fraction of solvent exposed Trp residues and *vice versa*. Therefore, for an unfolding protein one expects the *A*/*R* to decrease, but the CSM_0_ to increase, compared to a stabilised protein where one expects the *A*/*R* to decrease but the CSM_0_ to decrease. We note that a mixture of scenarios is possible and likely, e.g. the presence of both unfolded and aggregated material. However, our data demonstrate that the QUBES data show remarkable power and discriminatory ability.

Using this approach, it is possible to accurately detect, separate and quantify both protein unfolding and early stage formation of soluble aggregates as well as a predictor of sample stability. Our data suggest that the reason the approach is so sensitive is because it is based on the detection of a proteins intrinsic dynamic profile, which itself is a metric of changes to the proteins FEL and molecular flexibility. However, it is important to note that this rationale is a working hypothesis. Using QUBES has the advantage that: (i) data acquisition and analysis is rapid (<5 mins) so can be used as part of large scale screening; (ii) it can be used with any protein which includes one or more Trp residues (most proteins), (iii) using proteins of any size and in nearly any solvation/buffer environment; (iv) broad range of sample concentrations (*μ*g - mg) (v) samples are not consumed. We would stress that the approach is potentially useful in a comparative fashion and will be most robust and find best utility when examining e.g. variants of buffer conditions for the same protein. Moreover, it is difficult to envisage the approach being used with polyclonal antibodies where normal heterogeneity will be very large. That is, we expect the approach will be limited in use to homogenously purified single proteins.

## Author Contributions

MK, REW, AK, SP, and OK performed experiments. PP, HBLJ and SAW performed computational work. MC, AW, JMHE, AT, JO and CRP designed experiments and wrote the manuscript text. All authors contributed to data interpretation and analysis.

## Acknowledgements

The authors would like to thank Dr David Hough and Professor Randall Mrsny for informative discussions. We thank Bath ASU for the kind contribution of the commercially available antibodies used in the study.

## Supporting Information Available

Calculated mAb Fab tryptophan data and structure comparison; validating DLS profiles; results of structure-based calculations.

## Notes

We declare that CRP has commercial interests relating to the work presented in the manuscript, pertaining to a directorship of Bloc Laboratories Limited.

## References

1. Henzler-Wildman, K., and Kern, D. Dynamic personalities of proteins. Nature. 2007;450, 964–972. DOI: 10.1038/nature06522.

2. Wang, Z., Singh, P., Czekster, C. M., Kohen, A., and Schramm, V. L. Protein mass-modulated effects in the catalytic mechanism of dihydrofolate reductase: Beyond promoting vibrations. J. Am. Chem. Soc. 2014;136, 8333–8341. DOI: 10.1021/ja501936d.

3. Luk, L. Y. P., Ruiz-Pernía, J. J., Dawson, W. M., Roca, M., Loveridge, E. J., Glowacki, D. R., Harvey, J. N., Mulholland, A. J., Tuñón, I., Moliner, V., and Allemann, R. K. Unraveling the role of protein dynamics in dihydrofolate reductase catalysis. Proc. Natl. Acad. Sci. 2013;110, 16344–16349.

4. Pudney, C. R., Guerriero, A., Baxter, N. J., Johannissen, L. O., Waltho, J. P., Hay, S., and Scrutton, N. S. Fast protein motions are coupled to enzyme H-transfer reactions. J. Am. Chem. S.oc. 2013; 135, 2512–2517. DOI: 10.1073/pnas.1312437110

5. Klinman, J. P., and Kohen, A. Hydrogen Tunneling Links Protein Dynamics to Enzyme Catalysis. Annu. Rev. Biochem. 2013;82, 471–496. DOI: 10.1146/annurev-biochem-051710-133623.

6. Mauldin, R. V., Carroll, M. J., and Lee, A. L. Dynamic Dysfunction in Dihydrofolate Reductase Results from Antifolate Drug Binding: Modulation of Dynamics within a Structural State. Structure. 2009;17, 386–394. DOI: 10.1016/j.str.2009.01.005.

7. Zwiderek, K., Tunon, I., Moliner, V. & Bertran, J. Protein Flexibility and Preorganization in the Design of Enzymes. The Kemp Elimination Catalyzed by HG3.17. ACS Catal. 2015;5, 2587–2595. DOI: 10.1021/cs501904w.

8. varez-Garcia, D., and Barril, X. (2014) Relationship between Protein Flexibility and Binding: Lessons for Structure-Based Drug Design. J. Chem. Theory Comput. 2014;10, 2608–2614. DOI: 10.1021/ct500182z

9. Adhikary, R., Yu, W., Oda, M., Zimmermann, J., and Romesberg, F. E. Protein dynamics and the diversity of an antibody response. J. Biol. Chem. 2012;287, 27139–27147.

10. Thielges, M. C., Zimmermann, J., Yu, W., Oda, M., and Romesberg, F. E. Exploring the energy landscape of antibody-antigen complexes: Protein dynamics, flexibility, and molecular recognition. Biochemistry. 2008;47, 7237–7247. DOI: 10.1074/jbc.M112.372698.

11. Balakrishnan S. Moorthy, Steven G. Schultz, Sherry G. Kim, and Elizabeth M. Topp. Predicting Protein Aggregation during Storage in Lyophilized Solids Using Solid State Amide Hydrogen/Deuterium Exchange with Mass Spectrometric Analysis (ssHDX-MS). Mol. Pharmaceut. 2014;11, 1869–1879. DOI: 10.1021/mp500005v

12. Elvin, J. G., Couston, R. G., and Van Der Walle, C. F. (2013) Therapeutic antibodies: Market considerations, disease targets and bioprocessing. Int. J. Pharm. 2013;440, 83–98. DOI: 10.1016/j.ijpharm.2011.12.039.

13. Arbogast, L. W., Brinson, R. G., and Marino, J. P. Mapping Monoclonal Antibody Structure by 2D 13C NMR at Natural Abundance. Anal. Chem. 2015;87, 3556–3561. DOI: 10.1021/ac504804m.

14. Sobolewska-Stawiarz, A., Leferink, N. G. H., Fisher, K., Heyes, D. J., Hay, S., Rigby, S. E. J., and Scrutton, N. S. Energy landscapes and catalysis in nitric-oxide synthase. J. Biol. Chem. 2014;289, 11725–11738. DOI: 10.1074/jbc.M114.548834.

15. Laursen, T., Singha, A., Rantzau, N., Tutkus, M., Borch, J., Hedegård, P., Stamou, D., Møller, B. L., and Hatzakis, N. S. Single molecule activity measurements of cytochrome P450 oxidoreductase reveal the existence of two discrete functional states. ACS Chem. Biol. 2014;9, 630–634. DOI: 10.1021/cb400708v.

16. Ferreon, A. C. M., Ferreon, J. C., Wright, P. E., and Deniz, A. a. Modulation of allostery by protein intrinsic disorder. Nature. 2013;498, 390–394. DOI: 10.1038/nature12294.

17. Lanucara, F., Holman, S. W., Gray, C. J., and Eyers, C. E. The power of ion mobility-mass spectrometry for structural characterization and the study of conformational dynamics. Nat. Chem. 2014;6, 281–94. DOI: 10.1038/nchem.1889.

18. Zhang, A., Fang, J., Chou, R. Y.-T., Bondarenko, P. V., and Zhang, Z. Conformational Difference in Human IgG2 Disulfide Isoforms Revealed by Hydrogen/Deuterium Exchange Mass Spectrometry. Biochemistry. 2015;54, 1956–1962. DOI: 10.1021/bi5015216.

19. Tsai, C. J., Buyong, M., Sham, Y. Y., Kumar, S., and Nussinov, R. Structured disorder and conformational selection. Proteins Struct. Funct. Genet. 2001;44, 418–427. DOI: 10.1002/prot.1107.

20. Milanesi, L., Waltho, J. P., Hunter, C. a., Shaw, D. J., Beddard, G. S., Reid, G. D., Dev, S., and Volk, M. Measurement of energy landscape roughness of folded and unfolded proteins. Proc. Natl. Acad. Sci. 2012;109, 19563–19568. DOI: 10.1073/pnas.1211764109.

21. Reshetnyak, Y. K., Koshevnik, Y., and Burstein, E. A. Decomposition of Protein Tryptophan Fluorescence Spectra into Log-Normal Components. III. Correlation between Fluorescence and Microenvironment Parameters of Individual Tryptophan Residues. Biophys. J. 2001;81, 1735–1758. DOI: 10.1016/S0006-3495(01)75825-0

22. Muiño, P. L., and Callis, P. R. Solvent effects on the fluorescence quenching of tryptophan by amides via electron transfer. Experimental and computational studies. J. Phys. Chem. B. 2009;113, 2572–2577. DOI: 10.1021/jp711513b.

23. Moens, P. D. J., Helms, M. K., and Jameson, D. M. Detection of tryptophan to tryptophan energy transfer in proteins. Protein J. 2004;23, 79–83. DOI: 10.1023/B:JOPC.0000016261.97474.2e.

24. Demchenko AP. The red-edge effects: 30 years of exploration. Luminescence. 2002;17,19–42. DOI: 10.1002/bio.671

25. Chattopadhyay A & Haldar S. Dynamic insight into protein structure utilizing red edge excitation shift. Acc. Chem. Res. 2014;47, 12–19. DOI: 10.1021/ar400006z.

26. Tohru Azumi and Ken-ichi Itoh. Shift of emission band upon excitation at the long wavelength absorption edge. 1. A preliminary survey for quinine and related compounds. Chem. Phys. Lett. 1973;22, 395–399. DOI: 10.1016/0009-2614(73)80576-7.

27. Ken-ichi Itoh and Tohru Azumi. Shift of emission band upon excitation at the long wavelength absorptio edge. 2. Importance of the solute–solvent interaction and the solvent reorientation relaxation process. J Chem. Phys. 1975;62, 3431. DOI: 10.1063/1.430977.

28. Tohru Azumi, Ken-ichi Itoh, and Hiroshi Shiraishi. Shift of emission band upon the excitation at the long wavelength absorption edge. III. Temperature dependence of the shift and correlation with the time dependent spectral shift. J. Chem. Phys. 1976;65, 2550. DOI: 10.1063/1.433440.

29. Mitra M, Chaudhuri A & Patra M Organization and Dynamics of Tryptophan Residues in Brain Spectrin : Novel Insight into Conformational Flexibility. J Fluoresc. 2015;25, 707–17. DOI: 10.1007/s10895-015-1556-7.

30. Chakraborty, H, and Chattopadhyay, A. Sensing Tryptophan Microenvironment of Amyloid Protein Utilizing Wavelength-Selective Fluorescence Approach. J. Fluoresc. 2017;27, 1995–2000. DOI: 10.1007/s10895-017-2138-7.

31. Chattopadhyay A, Rawat SS, Kelkar D, Ray S & Chakrabarti A. Organization and dynamics of tryptophan residues in erythroid spectrin: novel structural features of denatured spectrin revealed by the wavelength-selective fluorescence approach. Protein Sci. 2003;12, 2389–2403. DOI: 10.1110/ps.03302003.

32. Rawat S S, Kelkar D & Chattopadhyay A. Monitoring gramicidin conformations in membranes: a fluorescence approach. Biophys. J. 2004;87, 831–843. DOI: 10.1529/biophysj.104.041715

33. Kelkar D, Chaudhuri A, Haldar S & Chattopadhyay A. Exploring tryptophan dynamics in acid-induced molten globule state of bovine alpha-lactalbumin: a wavelength-selective fluorescence approach. Eur. Biophys. J. 2010;39, 1453–1463. DOI: 10.1007/s00249-010-0603-1.

34. Catici DAM, Amos HE, Yang Y, van den Elsen JMH, Pudney CR. The red edge excitation shift phenomenon can be used to unmask protein structural ensembles: implications for NEMO–ubiquitin interactions. FEBS J. 2016;283, 2272–2284. DOI: 10.1111/febs.13724.

35. Jones HBL, Wells SA, Prentice EJ, Kwok A, Liang LL, Arcus VL, Pudney CR. A complete thermodynamic analysis of enzyme turnover links the free energy landscape to enzyme catalysis. FEBS J. 2017;284, 2829–42. DOI: 10.1111/febs.14152.

36. Gulácsy, C.E., Meade, R., Catici, D. A. M., Soeller, C., Pantos, G. D., Jones, D. D., Alibhai, D., Jepson, M., Valev, V. K., Mason, J. M., Williams, R. J. & Pudney, C. R. Excitation-energy-dependent molecular beacon detects early stage neurotoxic aβ aggregates in the presence of cortical neurons. ACS Chem. Neurosci. 2018; 10,1240–1250. DOI: 10.1021/acschemneuro.8b00322. DOI: 10.1021/acschemneuro.8b00322.

37. Jacobs, D. J., Rader, a. J., Kuhn, L. a, and Thorpe, M. F. Protein flexibility predictions using graph theory. Proteins Struct. Funct. Bioinforma. 2001;44, 150–165. DOI: 10.1002/prot.1081.

38. Wells, S. A., Jimenez-Roldan, J. E., Roemer, R. A., and Romer, R. A. Comparative analysis of rigidity across protein families. Phys. Biol. 2009;6, 46005–46011. DOI: 10.1088/1478-3975/6/4/046005.

39. Wells, S. a., Crennell, S. J., and Danson, M. J. Structures of mesophilic and extremophilic citrate synthases reveal rigidity and flexibility for function. Proteins Struct. Funct. Bioinforma. 2014;2657–2670. DOI: 10.1002/prot.24630.

40. Boning Liu, Huaizu Guo, Jin Xu, Ting Qin, Lu Xu, Junjie Zhang, Qingcheng Guo, Dapeng Zhang, Weizhu Qian, Bohua Li, Jianxin Dai, Sheng Hou, Yajun Guo & Hao Wang. Acid-induced aggregation propensity of nivolumab is dependent on the Fc. mAbs. 2015;8,1107–1117. DOI: 10.1080/19420862.2016.1197443.

41. Daniel, R. M., Danson, M. J., Eisenthal, R., Lee, C. K., and Peterson, M. E. The effect of temperature on enzyme activity: New insights and their implications. Extremophiles. 2008;12, 51–59. DOI: 10.1007/s00792-007-0089-7.

42. Morais, M. P. P., Fossey, J. S., James, T. D., and Van Den Elsen, J. M. H. Analysis of protein glycation using phenylboronate acrylamide gel electrophoresis. Methods Mol. Biol. 2012;869, 93–109. DOI: 10.1007/978-1-4939-8793-1_16.

