## Supplemental tables and figures for "Monoclonal antibody stability can be usefully monitored using the excitation-energy-dependent fluorescence edge-shift"

**Table S1. Summary of calculated parameters for mAb Fab regions**

| | Number of Trp residues in the Fab | Average Trp SASA trp (Å$^2$) | Average normalised Trp B-factor | Cumulative frequency of calculated normal modes | All atom flexible motion SVRC |
|---|---|---|---|---|---|
| | | | **Chimeric** | | |
| Cetuximab | 6 | 31.7 | 0.583 | 456.4 | 0.60 |
| Infliximab | 6 | 29.3 | 0.630 | 444.5 | 0.44 |
| Rituximab | 5 | 8.8 | 0.648 | 394.2 | 0.55 |
| | | | **Humanised** | | |
| Bevacizumab | 7 | 30.3 | 0.528 | 454.5 | 0.67 |
| Natalizumab | 5 | 8.9 | 0.620 | 392.4 | 0.50 |
| Pertuzumab | 4 | 2.6 | 0.510 | 502.3 | 0.75 |
| Trastuzumab | 5 | 7.4 | 0.639 | 435.8 | 0.56 |

**Figures**

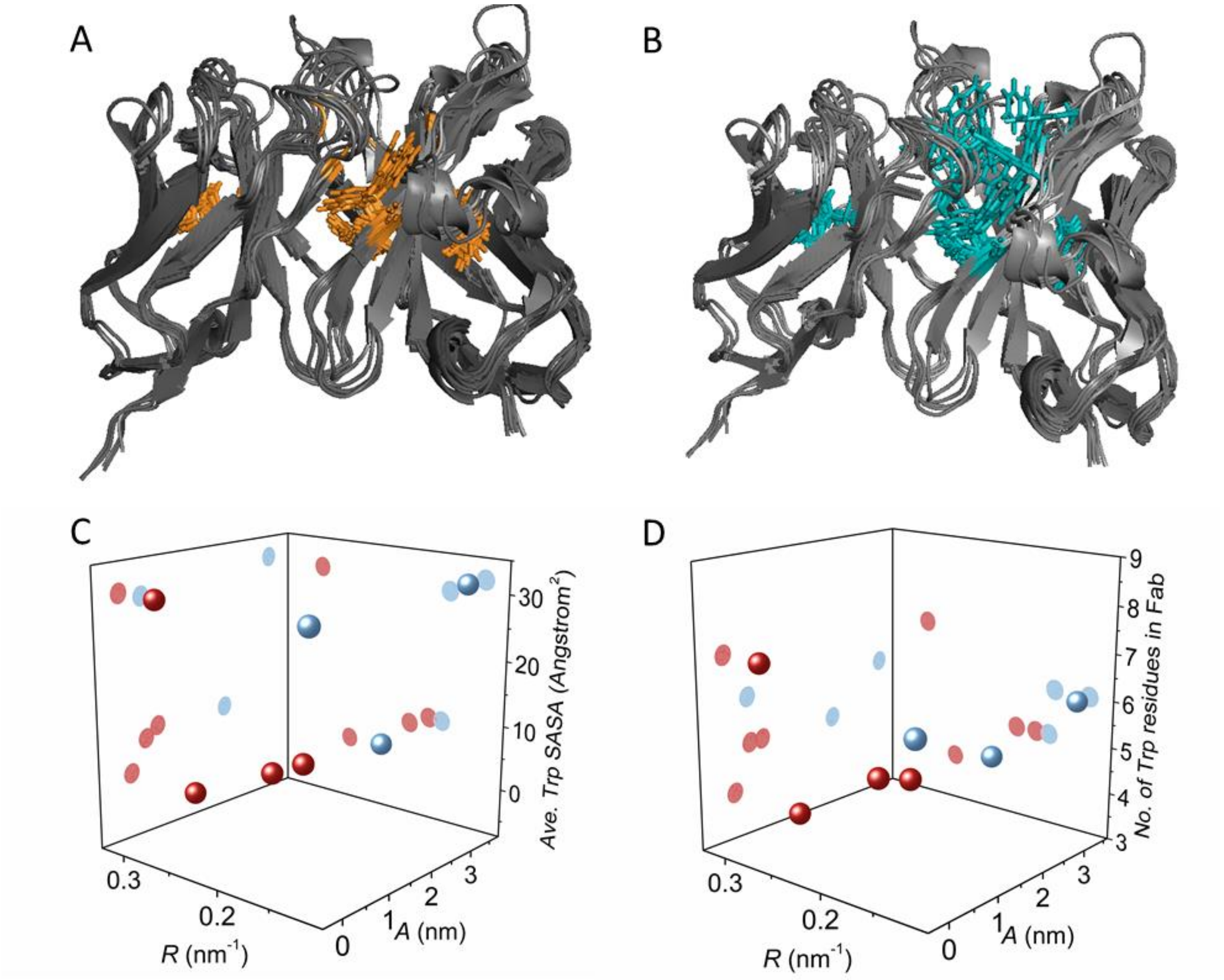


**Figure S1.** A PyMol representation of an overlay of all RosettaAntibody homology models. **A,** The conserved tryptophans have been displayed in stick representation and are highlighted in orange. It can be seen that the conserved 47th residues on the $V_H$ chain (boxed) have been modelled at two different orientations. The other conserved tryptophan residues are modelled in the same orientation. **B,** The same image, but with all (conserved and non-conserved) residues highlighted in blue. It is clear that the additional un-conserved tryptophan residues are located in a similar area of the molecule, and appear to be closer to the edge of the molecule (less-buried). Within each mAb Fab studied, there are between 4-7 tryptophan residues. The majority of these residues are conserved in the same position within the framework regions of the chains and are presented in the same orientation. The non-conserved tryptophans (Figure S1B that are less buried, reflected in elevated SASA values. Some of these residues are found in the CDR loops. **C,** Correlation of the quantified REES values with Trp solvent accessible surface area (SASA) in the Fab region. **D,** Correlation of the quantified REES values with the number of Trp residues in the Fab region.

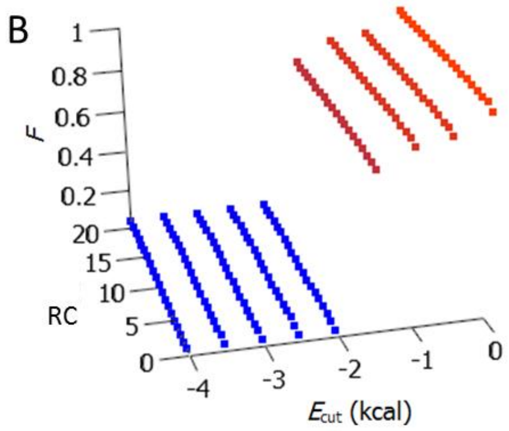


**Figure S2.** Example results from the rigid cluster decomposition analysis showing how the fraction of residues (*F*) in rigid clusters (*RC*) varies with energy cutoff ($E_{cut}$). The average of *E*cut value (sum value of rigid clusters; SVRC) is then used to quantify the difference in calculated flexibility between the mAbs.

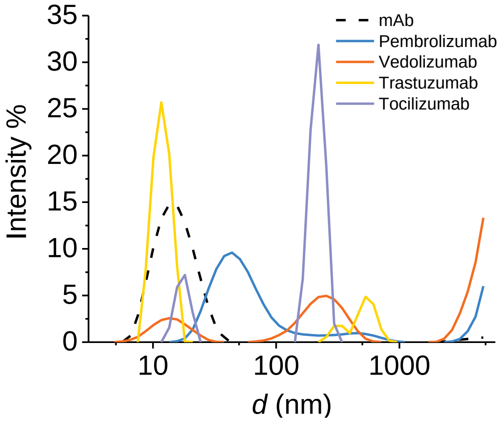


**Figure S3.** DLS profiles for thermally aggregated mAbs shown in Figure 4A.
